# Model-guided metabolic engineering of curcuminoid Production in *Pseudomonas putida*

**DOI:** 10.1101/2024.02.08.579459

**Authors:** Maria Martin-Pascual, Sara Moreno-Paz, Rik P. van Rosmalen, Julia Dorigo, Francesca Demaria, Richard van Kranenburg, Vitor A.P. Martins dos Santos, Maria Suarez-Diez

## Abstract

Production of value-added, plant-derived compounds in microbes increasingly attracts commercially interest in food and pharmaceutical industries. However, plant metabolic pathways are complex, require a robust balance of enzymes, cofactors, ATP and other metabolites, and often result in low production when transplanted to bacteria. This is exemplified by the biosynthesis of curcuminoids from the *Curcuma longa* plant. Here, we combine dynamic pathway modeling, systematic testing of isoenzymes, and the optimization of gene expression levels and substrate concentrations for the biosynthesis of curcuminoids in *Pseudomonas putida*, leading to unprecedented conversion rates of caffeic acid and tyrosine to curcumin. The development of kinetic ensemble models guided the design of production strains, emphasizing the necessity of high relative expression of *c3h, curs2* and *dcs* and, the low relative expression of *tal, comt, ccoaomt*, and *4cl4*. This optimization resulted in a strain that achieved a 10.8 ±1.8% of the maximum theoretical yield of curcumin from tyrosine. This represents a 4.1-fold increase in production efficiency and the highest yield reported to date, demonstrating the potential of *P. putida* as a promising platform for curcuminoid production. Our findings highlight the effectiveness of our strategy not only in the advances in the production of curcuminoids but also in setting a framework for the biosynthesis of other complex compounds.

## Introduction

Curcuminoids are polyphenolic compounds naturally found in the rhizome of the *Curcuma longa* (turmeric) plant. They account for 1 to 6 % (w/w) of turmeric rhizome, with curcumin being the most abundant, constituting 60 to 70 % (w/w) of the total curcuminoids, followed by demethoxycurcumin (20-27 % w/w) and bisdemethoxycurcumin (10-15 % w/w)^1^. Due to their characteristic strong yellow color, curcuminoids are widely used as coloring agents in the food industry and have been authorized as food additive by the European Union (code E100) and the FDA (label 73.615)^2,3^. Besides their use in the food industry, the pharmaceutical and cosmetic industries account for 50% of the curcuminoids global market^4^. Multiple biological activities, predominantly anti-carcinogenic, antioxidant and anti-inflammatory, have been attributed to curcumin and its derivatives, however, due to their poor pharmacokinetic properties, further research is needed to ascertain their full therapeutic potential^1,5^.

The curcumin global market size was over USD 58 million in 2020, and it is predicted to grow at a compound annual growth rate (CAGR) of 16.1 % by 2028^4^. Meeting such global demands with plant-based curcumin production is challenging due to the seasonal dependent growth of the turmeric plant which is hindered by the expected raise of temperatures^6^. Furthermore, traditional methods for the extraction of curcuminoids from the rhizome of the plant are lengthy, laborious, and require the use of organic solvents, high pressure and elevated temperatures. Although alternative methods hold promise for green extractions, they are inefficient, only able to extract up to 6% of the curcuminoids present in the rhizome^7^. Chemical synthesis of curcuminoids also has severe downsides, including the reliance on fossil fuel-derived solvents, toxic reagents and expensive starting compounds^8^. Microbial production of curcuminoids provides a valuable alternative to these processes, as it has the potential to provide a greater level of control, consistency, and efficiency, whilst operating at milder conditions and relying on biomass-derived feedstocks such as glucose, tyrosine, ferulic, p-coumaric and caffeic acids. This allows, in principle, for efficient, biobased and standardized large-scale production of curcuminoids with the potential to meet the increasing global demands.

The curcuminoid pathway has been expressed in various microorganisms, with a strong focus on *E. coli*, where the maximum reported yields have been obtained (Table 1). The conversion of ferulic acid to curcumin requires only three catalytic reactions, and 100% yields have been achieved^9^. Although the same enzymatic steps are required for production from p-coumaric acid, the maximum reported yield is 59% of the maximum theoretical yield^10^. When caffeic acid or tyrosine are used as substrates, the highest curcumin yields decrease to 2.12 and 1.27 % of the maximum theoretical yield, respectively^10,11^. These lower yields can be caused by the increased complexity of the pathway or the higher toxicity of these substrates and results in not yet commercially viable bio-based production of curcuminoids.

**Table 1:** Summary of microbial curcuminoids production. FEA, ferulic acid; CUA; p-coumaric acid; CAA, caffeic acid; TYR, tyrosine; CUR, curcumin; BDC, bisdemethoxycurcumin. The yield is expressed as a percentage of the maximum theoretical yield.

| Sub. | Organism | Genes | %-Yield | Ref. |
| --- | --- | --- | --- | --- |
| FEA | *E. coli* (1 mM) | *4cl, accABCD, cus* | CUR (61%) | 10 |
|  | *E. coli* (2 mM) | *4cl, dcs, curs1* | CUR (96%) | 20 |
|  | *E. coli* (2 mM) | *dcs, curs1, tal, c3h, ccoaomt,* | CUR (19%) | 11 |
|  | *E. coli* (2 mM) | *4cl tal, c3h, 4cl1, dcs, curs1* | CUR (91.7%) | 21 |
|  | *E. coli* (3 mM) | *tal, c3h, 4cl, comt, dcs, curs1* | CUR (100%) | 9 |
|  | *E. coli* (2 mM) | *cus, 4cl* | CUR (1.5%) | 22 |
| | *E. coli* $\Delta curA$ (4 mM) | *cus, 4cl*, improve membrane & MalCoA | CUR (73%) | 23 |
|  | *S. cerevisiae* (0.08 mM) | *4cl, ferA, cus, dcs, curs1* | CUR (17.8%) | 24 |
| CUA | *E. coli* (1 mM) | *4cl, accABCD, cus cus,* | BDC ( 59%) | 10 |
|  | *E. coli* (0.15 mM) | *4cl, accABCD, matBC, tal* | BDC (58.6%) | 25 |
| | *P. putida* $\Delta ech$ (5 mM) | *cus* | BDC (0.2%) | 26 |
|  | *E. coli* (2 mM) | *dcs, curs1, tal, c3h, ccoaomt, 4cl* | BDC (0.08%) | 11 |
|  | *Y. lipolytica* (2 mM) | *tal, cus, 4cl, g2ps1, pks1, sts, bas* | BDC (0.05%) | 27 |
| CAA | *E. coli* (1 mM) | *dcs, curs1, tal, c3h, ccoaomt, 4cl* | CUR (2.12%) | 11 |
| TYR | *E. coli* (3 mM) | *pal, 4cl, accABCD, cus* | BCUR (11.5%) | 10 |
|  | *E. coli* (3 mM) | *dcs, curs1, tal, c3h, ccoaomt, 4cl* | CUR (0.04%) | 11 |
|  | *E. coli* (3 mM) | *tal, c3h, 4cl, comt, dcs, curs1* | CUR (1.27%) | 9 |
|  | *E. coli* (3 mM) | *tal, 4cl4, cus* | BCUR (6.2%) | 28 |
|  | *E. coli* (3 mM) | *tal, 4cl4, cus, c3h, comt* | CUR (0.1%) | 28 |

The curcuminoids biosynthetic pathway has been reconstructed based on curcuminoid production in *C. longa* and *Oryza sativa*, and expanded to include enzymatic reactions derived from various yeast, bacterial and plant species (Figure 1)^10,12^. Curcuminoids can be formed from two l-tyrosine molecules in several steps. First, tyrosine is deaminated to form p-coumaric acid by tyrosine ammonia lyase (TAL). Next, p-coumaric acid can be converted to caffeic acid by coumarate-3-hydroxylase (C3H) or to coumaroyl-CoA by feruloyl/coumaroyl-CoA synthase (FCS) or 4-coumarate-CoA ligase (4CL). Subsequently, caffeic acid can be directly converted into ferulic acid by caffeic acid O-methyl transferase (COMT). Alternatively, it can be ligated to coenzyme-A (CoA) by FCS or 4CL, creating caffeoyl-CoA, which later can be converted to feruloyl-CoA by caffeoyl-CoA O-methyl transferase (CCOAOMT). When ferulic acid is formed, it can also be bound to CoA by FCS or 4CL, creating feruloyl-CoA. Hence, all three so-called hydroxycinnamic acids (ferulic, p-coumaric and caffeic acids) can be ligated to CoA by the action of either FCS or 4CL. Once the CoA-esters coumaroyl- and feruloyl-CoA are formed, malonyl-CoA is added to these compounds in a condensation reaction catalyzed by diketide-CoA synthase (DCS). Finally, the diketide-CoA esters react with a single molecule of a CoA ester, coumaroyl-CoA or feruloyl-CoA, to form a curcuminoid. This last reaction step can be catalyzed by curcumin synthase (CURS). When feruloyl-diketide-CoA reacts with a molecule of feruloyl-CoA, curcumin is formed. When coumaroyl-diketide-CoA reacts with coumaroyl-CoA, bisdemethoxycurcumin is formed, and when either diketide-CoA reacts with one of the other CoA esters (i.e. feruloyl-diketide-CoA with coumaroyl-CoA or vice versa), the asymmetric demethoxycurcumin is formed. There are three subtypes of CURS which showed to have different substrate specificity; CURS1 and CURS2 preferred feruloyl-CoA as a starting substrate, while CURS3 did not show any preference between feruloyl-CoA and coumaroyl-CoA^12^. Furthermore, these last two reactions, catalyzed by DCS and CURS, can also be performed by a single enzyme, curcuminoid synthase (CUS) that, although has a preference for coumaroyl-CoA and the production of bisdemethoxycurcumin, is also able to produce curcumin in trace amounts^10^.

**Figure 1:**
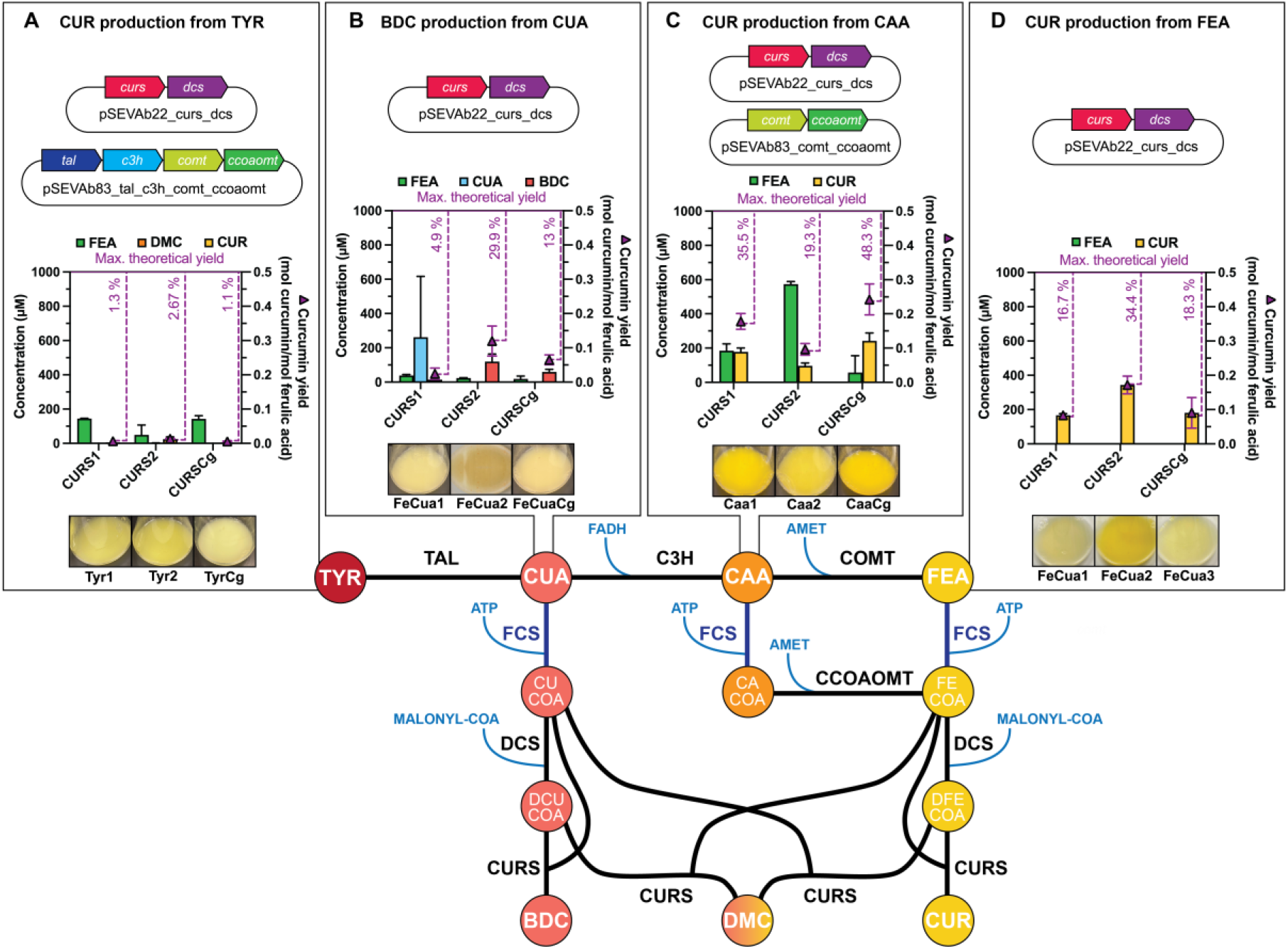
Overview of the curcuminoid production pathway showing the potential substrates and products. **A**. Curcumin (CUR) production from tyrosine (TYR), the performance of the three best strains is shown (see Sup. Figure 5 for additional information). **B**. Bisdemethoxycurcumin (BDC) production from p-coumaric acid (CUA), the performance of the three best strains is shown (see Sup. Figure 3 for additional information). **C**. Curcumin production from caffeic acid (CAA), the performance of the three best strains is shown (see Sup. Figure 4 for additional information). **D**. Curcumin production from ferulic acid (FEA), the performance of the three best strains is shown (see Sup. Figure 2 for additional information). Metabolite abbreviations: TYR, tyrosine; CUA, p-coumaric acid; CAA, caffeic acid; FEA, ferulic acid; CUCOA, coumaroyl-CoA; CACOA, caffeoyl-CoA; FECOA, feruloy-CoA; DCUCOA, diketide coumaroyl-CoA; DFECOA, diketide feruloyl-CoA; BDC, bisdemethoxycurcumin; DMC, demethoxycurcumin; CUR, curcumin; AMET, S-adenosyl-l-methionine. Enzyme abbreviations: TAL, tyrosine ammonia lyase; C3H, coumarate-3-hydroxylase; COMT, caffeic acid O-methyl transferase; FCS, feruloyl/ coumaroyl-CoA synthase; CCOAOMT, caffeoyl-CoA O-methyl transferase; DCS, diketide-CoA synthase; CURS, curcumin synthase.

**Figure 2:**
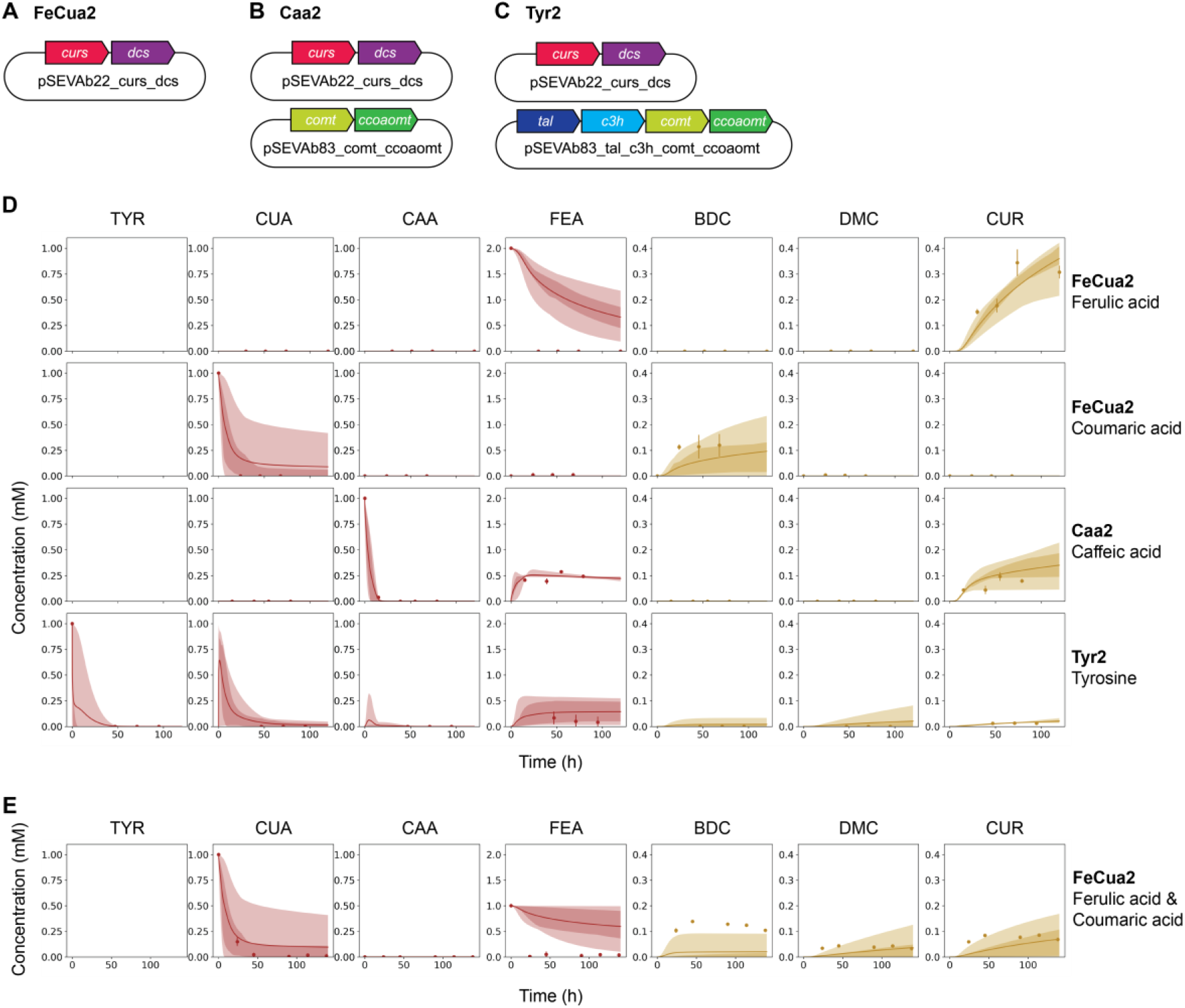
Comparison of ensemble model simulations and experimental data. **A-C**. Plasmids carried by the strains used for parameter estimation (FeCua2, Caa2 and Tyr2). **D**. Ensemble models fit to the experimental data used for parameter estimation, the strains and substrate used are indicated. **E**. Ensemble models fit to experimental data of validation experiment, the strain and substrates used is indicated.

The presence of promiscuous enzymes in the curcuminoid pathway complicates its optimization, as alterations in gene expressions can lead to unforeseen effects on reaction rates. To address this challenge, dynamic pathway models, which are constructed using a system of ordinary differential equations (ODEs), can be employed. These models account for metabolite concentrations and enzyme fluxes over time. They rely on mathematical representations of enzyme kinetics, such as Michaelis-Menten equations, and necessitate the determination of kinetic parameters^13,14^. They serve as a valuable tool for comprehending pathway fluxes, assessing (non-measurable) intermediate concentrations, and pinpointing potential limiting steps. Furthermore, they can be used to evaluate how alterations in enzyme concentrations impact production, thereby aiding in the design of optimized pathways^13,14^.

To enhance curcuminoid production, the choice of a microorganism able to endure toxic pathway metabolites is essential. *Pseudomonas putida* KT2440 has gained recognition as a promising platform for the production of various biological products. It offers a robust metabolism as well as a high tolerance to a variety of substances, particularly aromatic compounds^15^, making it a suitable candidate for tolerating the toxicity of the various substrates and intermediates in the curcuminoid pathway^15–18^. Natively, *P. putida* is able to catalyze the conversion of hydroxycinnamic acids into CoA ester molecules due to the presence of *fcs* (PP 3356). However, it is able to degrade the phenylpropanoidCoA metabolites due to the presence of *fcs* (PP 3356). However, it is able to degrade the phenylpropanoidCoA metabolites using *ech* (PP 3358), which interferes with the production of curcuminoids. Although production of bisdemethoxycurcumin from p-coumaric acid has been achieved in *P. putida* Δ*ech*, only low yields have been obtained^19^ and the potential of this host for curcuminoids production remains largely unexplored.

Here, we demonstrate the successful production of curcuminoids from ferulic acid, p-coumaric acid, caffeic acid and tyrosine facilitated by plasmid-based expression of heterologous genes in *P. putida* Δ*ech*. A curcumin yield of 34.4 ± 5.2 % of the maximum theoretical yield was reached from ferulic acid, 24.0 ± 8.7 % from p-coumaric acid, 48.5±9.1 % from caffeic acid and 2.7±0.2 % from tyrosine. Experimental data were used to create ensemble dynamic models of the curcuminoids pathway that connect enzyme concentrations, enzyme kinetics, thermodynamics and metabolite concentrations. These models served as a guide in designing new strains with varying expression levels of the pathway genes, resulting in the optimization of curcumin production from tyrosine up to a 10.8±1.8 % of the maximum theoretical yield, representing a 4.1-fold increase in production. This study highlights the potential of dynamic pathway models in comprehending intricate production pathways, providing a foundation for generating hypotheses that guide the optimization process. Moreover, the curcumin yields achieved from caffeic acid and tyrosine stand as the highest reported to date. Additionally, curcumin was successfully produced from glucose, showcasing the potential of *P. putida* as a promising cell factory for the production of curcuminoids.

## Material and methods

### Media, Bacterial strains and chemicals

Lysogeny-Broth (LB) (10 g/L tryptone, 10 g/L NaCl and 5 g/L yeast extract) and M9 minimal media ((1.63 g/L NaH2PO4, 3.88 g/L K2HPO4, 2 g/L(NH4)2SO4, 10 mg/L EDTA, 100 mg/L MgCl2•6H2O, 2 mg/L ZnSO4•7H2O, 1 mg/L CaCl2•2H2O, 5 mg/L FeSO4•7H2O, 0.2 mg/L Na2MoO4•2H2O, 0.2 mg/L CuSO4•5H2O, 0.4 mg/L CoCl2•6H2O, and 1 mg/L MnCl2•2H2O) were used to grow the bacterial strains. When necessary, antibiotics were added to the media: kanamycin (50 μ/ml), gentamycin (15 μ/ml), streptomycin (100 μ/ml), or chloramphenicol (34 μ/ml). *E. coli* λpir competent cells were prepared as described by Green and Rogers^29^ and were used for plasmid propagation. *E. coli* cells were grown in LB media at 37 °C in a shaking incubator at 200 rpm. *P. putida* cells were grown in LB and M9 media at 30 °C in a shaking incubator at 200 rpm. *P. putida* Δ*ech* strain was generated from *P. putida* KT2440 using the pGNW and pQURE plasmids^30^. *P. putida* Δ*ech* was made electrocompetent after several washing steps with 300 mM sucrose. A single exponential decay pulse was applied using the GenePulser XcellTM (Bio-Rad) set at 2500, 200 and 25 V.

A list of all strains, plasmids, and primers used in this study can be found in Supplementary Data 1. Analytical standards of l-tyrosine, ferulic acid, p-coumaric acid, caffeic acid, curcumin, demethoxycurcumin and bisdemethoxycurcumin were purchased from Sigma-Aldrich. Acetonitrile and hydrochloric acid were purchased from Acros Organics, and ethyl acetate and DMSO were purchased from Sigma-Aldrich.

### Toxicity essays

Toxicity assays were performed with all four precursors (ferulic, p-coumaric and caffeic acids, and tyrosine) on *P. putida* Δ*ech* in a concentration range of 0-0.25-0.5-1-2-4-8 mM. *P. putida* Δ*ech* was grown O/N at 30 °C in a 50 mL falcon tube containing 10 mL of LB. The next day, the O/N liquid culture was centrifuged for 10 min at 4700 g. The supernatant was discarded, and the cell pellet was resuspended in 10 mL of M9 media containing 70 mM of glucose. The OD600nm of the cultures was measured in a spectrophotometer (IMPLEN Westburg). A 96-well plate was prepared by adding 200 μL of the sample per well with a starting OD600nm of 0.2. The OD600nm readings were monitored in a microplate reader (BioTek Synergy/neo2 or BioTek SynergyMx) every 5 minutes for 48 hours at 30 °C with a continuous shake.

### Plasmid construction

Gene sequences of *tal, c3h, ccoaomt, dcs, curs1*, and *comt* were obtained from Rodrigues and colleagues^11^. Gene sequences of *curs2* and *curs3* were obtained from NCBI with the accession numbers AB506762 and AB506763, respectively. All the gene sequences were codon optimized for *P. putida* KT2440 using the Jcat codon optimization tool (www.jcat.de), and genes were ordered from Twist Bioscience. The genes for *curs1* and *dcs* codon optimized for *C. glutamicum* were kindly provided by dr. Melanie Mindt. Gene sequences can be found in Supplementary Data 1.

The genes were first ligated into the pSB1C3 repository plasmid via golden gate cloning using BsaI restriction enzyme. The BsaI overhangs were already present on the genes codon optimized for *P. putida*. The *curs* and *dcs* genes codon optimized for *C. glutamicum* were PCR amplified with M526M27 and M528-M529 pair of primers, respectively, using NEB Q5® High-Fidelity DNA polymerase according to manufacturer’s protocol (M0491). Next, PCR fragments were loaded onto 1 % agarose gel and purified using the NucleospinTM Gel and PCR Clean-up kit from Fischer ScientificTM. Plasmids were built using the SevaBrick Assembly method^31^ and transformed into *E. coli* αλpir competent cells by heat-shock. Colonies were screened for the correct assembly of the plasmid by Colony PCR using Phire Green Hot Start II polymerase. Colonies with the right size band were grown overnight in 10 mL of LB supplemented with the appropriated antibiotic. Glycerol stocks were prepared for longterm storage. Plasmids were purified from the overnight liquid cultures using the GeneJET Plasmid Miniprep kit from Thermo ScientificTM. Plasmid sequence was confirmed by Sanger sequencing from Macrogen (MACROGEN Inc. DNA Sequencing Service; Amsterdam, The Netherlands).

The synthetic pathway to produce curcuminoids was divided in three levels. Each level corresponds to a specific intermediate that serves as the starting substrate for the synthesis of curcuminoids: either ferulic or coumaric acid, caffeic acid or tyrosine. The first constructs were designed to convert ferulic or coumaric acid to curcumin. One of the different *curs* isoenzymes (1, 2 and 3 codon optimized for *P. putida* and type 1 codon optimized for *C. glutamicum*) and *dcs* (codon optimized for *P. putida*) were ligated into the pSEVAb22 backbone carrying a p100 promoter. In addition, the *curs1* gene codon optimized for *C. glutamicum* (*cursCg*) was ligated to *dcs* gene codon optimized for *C. glutamicum* (*dcsCg*) into the pSEVAb22 backbone. For the second construct, designed to convert caffeic acid to ferulic acid or feruloyl-coA, the *comt* and *ccoaomt* genes were ligated into pSEVAb83 backbone, carrying a p100 promoter. For the third construct, designed to convert tyrosine into coumaric acid, the *comt, ccoaomt, c3h* and *tal* genes were ligated into pSEVAb83, also carrying a p100 promoter. Additionally, different combinations of genes and plasmid backbones were constructed. The list of all plasmids and strains constructed is listed in Supplementary Data2.

### Curcuminoid production experiments

The different strains were grown in 10 mL LB medium containing the appropriate antibiotic, at 30 °C and 200 rpm, overnight. The next day, the overnight liquid cultures were centrifuged for 10 min at 4700 g. The supernatant was discarded and the cell pellet was washed in 1 mL of M9 media containing 70 mM glucoseduring the LB-to-M9 transition in order to eliminate LB traces. Cells were resuspended to an starting OD600nm of 0.3 in a total volume of 25 mL of fresh minimal M9 media supplemented with 70 mM of glucose, the appropriate antibiotics, and the precursor (ferulic -, p-coumaric -, caffeic acid or tyrosine) at indicated concentrations. The 250 mL-Erlenmeyer flasks, containing the cells were grown aerobically at 30°C and 200 rpm for 72 h. Samples were taken for each flask at indicated time points and phenylpropanoids and cucuminoids were extracted. Biological triplicates were included.

### Extraction of phenylpropanoids and curcuminoids

To measure curcuminoids and precursors, a sample of 500 μL (extraction from experiments with ferulic, coumaric or caffeic acid as precursors) or 1 mL (extraction from experiments with tyrosine as precursor, or a combination of ferulic and coumaric acid) was taken from each Erlenmeyer, and transferred to a clean 2 mL Eppendorf tube. Then, 1 μL of 6M HCl was added to each tube, and the tubes were vortexed to break the cells. To extract the curcuminoids, an equal amount of ethyl acetate was added to the Eppendorf. The tubes were then incubated at 55 °C for 10 minutes at 800 rpm. Then, the tubes were centrifuged in a microcentrifuge at 20238 g for 2 min. The top layer was transferred to a new tube, making sure none of the cell pellet was taken in the process. This extraction method was repeated until there was no yellow color visible in the cell pellet. The ethyl acetate was then evaporated in a rotary evaporator (Concentrator plus, Eppendorf) at 60°C. Finally, the remaining dry sample was dissolved in 500 μL of DMSO.

### Quantification of phenylpropanoids and curcuminoids

High-performance liquid chromatography (HPLC) was used to quantify the substrates (ferulic, pcoumaric, caffeic acid and tyrosine) and curcuminoids (demethoxycurcumin, bisdemethoxycurcumin and curcumin). HPLC was performed on a Shimadzu LC2030C machine equipped with a Poroshell 120EC-C18 column (250 x 4.6 mm, Agilent) and a UV/vis detector. Mobile phase A was composed of Milli-Q water, mobile phase B of 100 mM formic acid, and mobile phase C of acetonitrile. Mobile phase was used at a rate of 1 ml/min and was composed of Milli-Q water (A), 100 mM formic acid (B), and acetonitrile (C) at varying proportions: 77:10:13 (v/v/v) in the first 10 min, 23:10:67 (v/v/v) in the next 9 min, and 77:10:13 (v/v/v) in the last six minutes. Curcuminoids and hydroxycinnamic acids were detected at a wavelength of 420 nm and 280 nm, respectively.

Yields were calculated by dividing the maximum measured curcuminoid concentration by the consumed substrate. Yields are expressed as a percentage of the maximum curcuminoid yield (0.5 mol curcuminoid/ mol substrate).

### Construction of kinetic models of the curcuminoid pathway

Three models, mFeCua2, mCaa2, and mTyr2, corresponding to strains FeCua2, Caa2, and Tyr2, were developed based on the genotype of each strain (Table 2). For most reactions, the generalized reversible Michaelis-Menten kinetics expression rate law was applied. However, for reaction C3H, a mass-action rate law was used due to its high equilibrium constant^32^. The rate laws were formulated using the Wegscheider-compliant parametrization, which models the reaction equilibrium constant by considering the contributions of the concentration and chemical potential of each reactant. This approach helps avoid hidden dependencies between the equilibrium constants of multiple reactions, which could lead to thermodynamically infeasible parameterizations^32^. Enzymes catalyzing multiple reactions (FCS, DCS, and CURS) had additional substrate competition terms incorporated into their rate laws, as outlined in the appendix of Liebermeister et al.^32^. To simplify the model, the concentrations of all co-factors were set as constant values, but their chemical potential contributions to relevant rate laws were considered. Additionally, enzyme concentration and rate parameters were merged when possible to mitigate issues related to identifiability. Growth was integrated into the model through a logistic equation, which was used to scale enzyme and metabolite concentrations by the biomass concentration. The most complex model (mTyr2) encompassed 14 reactions and 13 ODEs to represent the dynamic concentrations of 12 metabolites and biomass. This model involved a total of 89 parameters, with 37 representing Michaelis-Menten constants (kM), 16 representing enzyme concentrations, rates, or a combination of both (u, kV, and uV, respectively), 23 representing standard thermodynamic potentials (μ), 9 representing fixed concentrations of co-factors (c), 2 being part of the logistic equation representing growth, and 2 being fixed (R and T). For detailed information regarding model construction and structure, please see the Supplementary Methods.

**Table 2:** Summary of constructed models.

| Model name | Included enzymes |
| --- | --- |
| mFeCua2 | FCS, DCS, CURS2 |
| mCaa2 | COMT, CCOAOMT, FCS, DCS, CURS2 |
| mTYR2 | TYR, C3H, COMT, CCOAOMT, FCS, DCS, CURS2 |

### Parameterization of kinetic models of the curcuminoid pathway

To establish initial parameter value ranges, various literature sources were combined. When available, co-factor concentration data was sourced from studies on *P. putida* KT2440^33^. In cases where data was not available, information was gathered from *Pseudomonas taiwanensis* VLB120^34^ and *E. coli* K12^35^, with preference given to the former. Estimates for chemical potentials were derived from changes in the Gibbs free energy of the reactions, computed using the eQuilibrator API under standard physiological conditions (pH = 7.5, ionic strength = 0.25 M, and pMg = 3)^36^. Enzyme kinetic parameters for DCS and CURS were obtained from Katsuyama et al.^37,38^. In instances of missing data, broad estimates were used. All parameter data was integrated through parameter balancing^39^, resulting in a final multivariate normal distribution for the parameters. The covariance^40^ between the estimated chemical potentials from eQuilibrator was included in the parameter balancing output, preserving the shared uncertainty in the estimates.

For parameter estimation of the mFeCua2, mCaa2, and mTyr2 models, experimental data included measurements of OD600nm and concentrations of substrates and curcuminoids from the FeCua2 strain grown in 2 mM ferulic acid or 1 mM p-coumaric acid, the Caa2 strain grown in 1 mM caffeic acid, and the Tyr2 strain grown in 1 mM tyrosine. Parameters were simultaneously estimated for all models using python (version 3.9.12) and the python libraries PESTO (0.2.12)^41^ and AMICI (version 0.11.21)^42^. Initial parameter estimates were sampled from the balanced distribution and log-transformed where appropriate (i.e., for parameters representing kinetic constants, enzyme concentrations, or co-factor concentrations). We employed multi-start local optimization (L-BFGS-B, 1000 starts, 100 maximum iterations), with a least-squares objective function that minimized the sum of squares of the residuals between the experimental data and the model predictions. Ensemble models were created for each strain (enFeCua2, enCaa2, and enTyr2), incorporating the top ten parameterized models. The goodness of fit for each observable (tyrosine, caffeic acid, p-coumaric acid, ferulic acid, bisdemethoxycurcumin, demethoxycurcumin, curcumin concentrations and OD600) in all experiments was evaluated using the mean square error normalized by the mean of the measurements. Simulations were performed using the AMICI library^42^ and the CVODES ODE solver^41^.

### Model-based optimization of curcumin production from tyrosine

Possible limiting reactions were identified following two complementary approaches. First, ensemble models were used to simulate the conducted experiments and, for each reaction, the maximum predicted flux was calculated. Reactions exhibiting lower flux values were flagged as potential bottlenecks, and the corresponding enzymes were selected as targets for over-expression. Furthermore, the enTyr2 ensemble was used to evaluate the influence of varying enzyme concentrations on curcumin production from tyrosine. The model was simulated under different enzyme concentration scenarios (ranging from 0 to 10 times the predicted concentration). Based on their impact on curcumin production, enzymes were considered as candidates for either over-expression or down-regulation. Expression of genes in plasmids with distinct copy numbers allowed the experimental implementation of model suggestions. pSEVAb22, pSEVAb83 and pSEVAb25 plasmids were used as low-copy-number, medium-copy-number and high-copy-number plasmids, respectively^31^.

### Code availability

The models in SBML format as well as the used parameters ranges can be found at https://gitlab.com/wurssb/Modelling/kinetics curcumin pathway. In addition, code for the simulation and analysis, as well as the required software packages and version information can be found in GitLab. Production and growth data of the performed experiments is available in Sup. Data 2.

## Results

### Curcuminoid substrate toxicity in *P. putida* Δ*ech*

To establish *P. putida* KT2440 as an efficient platform to produce curcuminoids, degradation of coumaroyl-CoA, caffeoyl-CoA and feruloyl-CoA by ECH should be avoided. Therefore, we deleted the *ech* gene and evaluated the toxicity of the four precursors (ferulic, p-coumaric and caffeic acids, as well as tyrosine) on *P. putida* Δ*ech*. In this way, the highest concentration of each precursor that does not inhibit growth was identified. The growth of *P. putida* Δ*ech* was hindered when ferulic and p-coumaric acids were present in the media at concentrations of 4 mM. Similarly, tyrosine at 8 mM and caffeic acid at 0.25 mM completely inhibited growth (Sup. Figure 1).

### Production of curcuminoids from ferulic and p-coumaric acid

Production of curcuminoids from ferulic and p-coumaric acids requires the expression of two heterologous genes, *dcs* and *curs*, and the native expression of *fcs* (Figure 1). Therefore, curcuminoid production from these substrates was considered as the first step towards the expression of the full curcuminoids pathway in *P. putida* Δ*ech*.

Five strains expressing *dcs* and *curs* in a pSEVAb22 plasmid backbone were constructed: FeCua1, FeCua2, FeCua3, FeCuaCg and FeCuaCgCg (Table 3). These strains differed on the *curs* isoenzyme used (*curs1, curs2* or *curs3*) and the codon optimization of *curs* and *dcs* (either for *P. putida* or *C. glutamicum* (Cg)). When these strains were grown in M9 supplemented with 1 mM and 2 mM of ferulic acid, they exhibited varied production levels (Table 3). A starting 2 mM concentration of ferulic acid resulted in the highest titers and curcumin production by FeCua2 (0.34 ± 0.05 mM), followed by FeCuaCg (0.18 ± 0.089 mM), FeCua1 (0.17 ± 0.01 mM) and FeCua3 (0.06 ± 0.02 mM) (Figure 1D, Sup. Figure 2). FeCuaCgCg grew slower and produced little amount of curcumin compared with the other strains (0.003 ±0.00 mM) (Sup. Figure 2). Although FeCua3 was the only strain that accumulated detectable levels of ferulic acid, the total conversion of ferulic acid to curcumin was below 100%, lower than the conversion rates previously reported in *E. coli* (Table 1)^43^. Moreover, increasing the initial OD600nm significantly influenced curcumin production. FeCua1 was inoculated at three different OD600nm, 0.3, 0.6 and 0.9 with 2 mM of ferulic acid. The highest titer, 0.60 ± 0.08 mM (60 % of the maximum theoretical yield), was achieved when ferulic acid was added at OD600nm 0.9 (Sup. Figure 2), representing a 0.74 fold-increase compared to the yield obtained with an initial OD600nm of 0.3.

**Table 3:** Production of curcumin from ferulic acid (FEA) or bisdemethoxycurcumin from coumaric acid (CUA). The mean and standard deviation of three replicates are shown. The yield is expressed as a percentage of the maximum theoretical yield. a indicates initial OD600nm = 0.6, b indicates initial OD600nm = 0.9, and all other production experiments were performed with an initial OD600nm = 0.3. p22 refers to pSEVAb22 plasmid backbone.

| Strain Name | Plasmids | Substrate (mM) | Titer (mM) | Yield (%) |
| --- | --- | --- | --- | --- |
| FeCua1 | p22-*curs1-dcs* | FEA (1 mM) | $0.05 \pm 0.02$ | $9.73 \pm 3.01$ |
| | | FEA (2 mM) | $0.17 \pm 0.01$ | $16.68 \pm 0.98$ |
| | | FEA (2 mM)<sup>a</sup> | $0.50 \pm 0.02$ | $50.00 \pm 1.56$ |
| | | FEA (2 mM)<sup>b</sup> | $0.64 \pm 0.03$ | $63.60 \pm 2.97$ |
| | | CUA(1 mM) | $0.01 \pm 0.00$ | $4.90 \pm 3.28$ |
| FeCua2 | p22-*curs2-dcs* | FEA (1 mM) | $0.16 \pm 0.02$ | $32.11 \pm 3.34$ |
| | | FEA (2 mM) | $0.34 \pm 0.05$ | $34.37 \pm 5.20$ |
| | | CUA (1 mM) | $0.12 \pm 0.04$ | $23.95 \pm 8.73$ |
| FeCua3 | p22-*curs3-dcs* | FEA (2 mM) | $0.06 \pm 0.00$ | $6.19 \pm 0.3$ |
| FeCuaCg | p22-*cursCg-dcs* | FEA (1 mM) | $0.07 \pm 0.01$ | $14.01 \pm 2.64$ |
| | | FEA (2 mM) | $0.18 \pm 0.09$ | $18.13 \pm 8.89$ |
| | | CUA (1 mM) | $0.06 \pm 0.01$ | $13.03 \pm 2.88$ |
| FeCuaCgCg | p22-*cursCg-dcsCg* | FEA (2 mM) | $0.00 \pm 0.00$ | $0.34 \pm 0.01$ |

FeCua1, FeCua2 and FeCuaCg were also grown in M9 media supplemented with 1 mM of p-coumaric acid (Figure 1 B, Table 3, Sup. Figure 3). Similar to the experiments with ferulic acid as substrate, FeCua2 produced the highest concentration of bisdemethoxycurcumin (0.12 ± 0.04 mM). Interestingly, small amounts of ferulic acid were detected in the samples, as well as trace amounts of demethoxycurcumin. This suggests that *P. putida* might synthesize ferulic acid from coumaric acid, even without expressing *c3h* and *comt* genes, which can lead to the production of demethoxycurcumin and curcumin from this substrate (Figure 1). Furthermore, although FeCua1 expresses *curs1* codon optimized for *P. putida* and FeCuaCg expresses *curs1* codon optimized for *C. glutamicum*, they differed on the produced bisdemethoxycurcumin (0.01 ± 0.00 mM and 0.06 ± 0.02 mM, respectively), showcasing the effect of codon optimization on production (Table 3). The highest conversion of p-coumaric acid was achieved by FeCua2 (24.0 ± 8.7%). Although this conversion is below the reported 60% conversions by *E. coli*^10,25^, it represents a considerable improvement to previous attempts of bisdemethoxicurcumin production by *P. putida* that showed conversions below 1% (Table 1)^26^.

**Figure 3:**
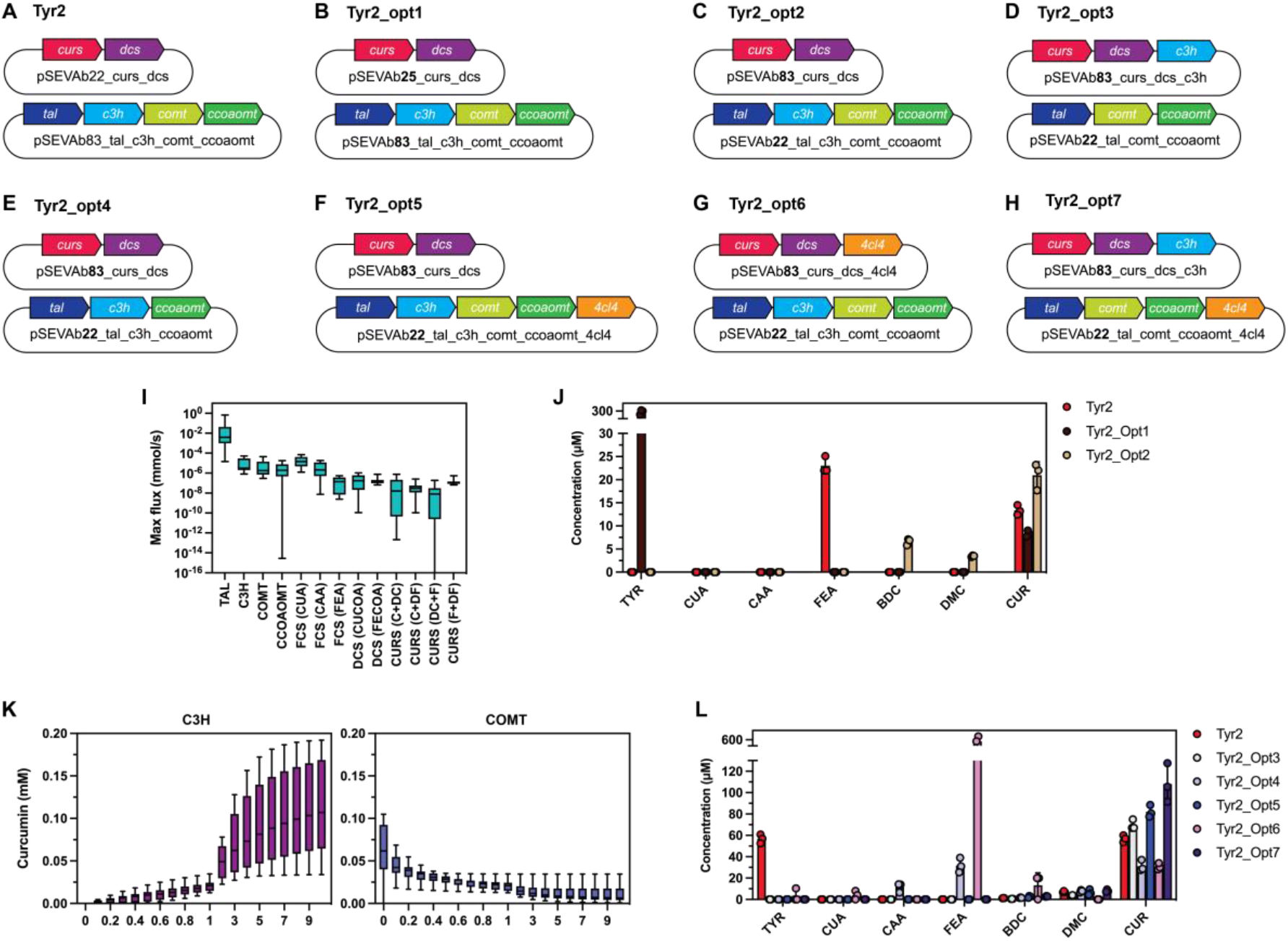
Model-based optimization of curcumin production from tyrosine. **A-H**. Plasmids carried by the optimized strains. **I**. Maximum predicted fluxes of each curcuminoid pathway reaction by enTyr2 ensemble. For promiscuous enzymes, the substrates of each reaction are indicated between brackets: pcoumaric acid (CUA, C), caffeic acid (CAA), ferulic acid (FEA, F), coumaroyl-CoA (CUCOA), feruloylCoA (FECOA), diketyde coumaroyl-CoA (DC), diketyde feruloyl-CoA (DF). **J**. Effect of changing relative expression of *dcs* and *curs2*. **K**. Predicted curcumin production based on relative C3H or COMT enzyme concentration predicted by enTyr2 ensemble. **L**. Effect of changing relative expression of *c3h* and *4cl4*, and omitting the expression of *comt*.

### Production of curcuminoids from caffeic acid

After the successful synthesis of curcuminoids from ferulic and p-coumaric acid, the biosynthetic pathway for curcuminoid production was further expanded to produce curcumin from caffeic acid. This involved the introduction of two additional genes into a pSEVAb83 plasmid: *comt* and *ccoaomt* (Figure 1). Through the expression of both the pSEVAb22-*curs*-*dcs* and the pSEVAb83-*comt*-*ccoaomt* plasmids, five *P. putida* Δ*ech* strains able to synthesized curcumin from caffeic acid were constructed: Caa1, Caa2, Caa3, CaaCg and CaaCgCg (Table 4).

**Table 4:** Production of curcumin from caffeic acid (CAA). The mean and standard deviation of three replicates are shown. The yield is expressed as a percentage of the maximum theoretical yield. p22 and p83 refer to pSEVAb22 and pSEVAb83 plasmid backbones, respectively.

| Name | Plasmids | CAA (mM) | Titer (mM) | Yield (%) |
| --- | --- | --- | --- | --- |
| **Caa1** | p22-*curs1-dcs* + p83-*comt-ccoaomt* | 1 | 0.17 ± 0.02 | 35.53 ± 4.56 |
| | | 2 | 0.00 ± 0.00 | 7.79 ± 0.45 |
| **Caa2** | p22-*curs2-dcs* + p83-*comt-ccoaomt* | 1 | 0.1 ± 0.02 | 19.27 ± 3.38 |
| | | 2 | 0.00 ± 0.00 | 0.13 ± 0.03 |
| **Caa3** | p22-*curs3-dcs* + p83-*comt-ccoaomt* | 2 | 0.00 ± 0.00 | 0.15 ± 0.04 |
| **CaaCg** | p22-*cursCg-dcs* + p83-*comt-ccoaomt* | 1 | 0.24 ± 0.05 | 48.47 ± 9.06 |
| | | 2 | 0.14 ± 0.01 | 14.84 ± 1.33 |
| **CaaCgCg** | p22-*cursCg-dcsCg* + p83-*comt-ccoaomt* | 2 | 0.00 ± 0.00 | 0.11 ± 0.19 |

Initially, the strains were tested with 2 mM caffeic acid. However, the curcumin levels were low (Caa1, Caa2, CaaCg) or zero (Caa3) and the lag phase of the cultures increased (Table 4, Sup. Figure 4), confirming the high toxicity of CAA found during the toxicity assay. Subsequently, Caa1, Caa2 and CaaCg, the strains with the best performance, were tested with 1 mM caffeic acid. The growth profile improved and the levels of curcumin significantly increased (Figure 1C, Table 4, Sup. Figure 4). Similar to the experiments with ferulic and p-coumaric acids, curcumin degradation was observed after 70 h. Caa1 and CaaCg produced 0.18 ± 0.02 and 0.24 ± 0.05 mM of curcumin, respectively while Caa2 produced 0.10 ± 0.02 mM. Furthermore, ferulic acid was accumulated in large amounts for Caa1 (0.19 ± 0.04 mM) and Caa2 (0.57 ± 0.02 mM). The highest caffeic acid conversion, achieved by CaaCg (48.47 ± 9.06%), was 22.8 times higher than that obtained using *E. coli* (Table 1)^11^.

### Production of curcuminoids form tyrosine

After successfully producing curcumin from caffeic acid, the curcuminoids pathway was expanded to include heterologous expression of *tal* and *c3h* (Figure 1). Four strains able to produce curcuminoids from tyrosine were built differing in the *curs* isoenzyme used and the codon optimization: Tyr1, Tyr2, Tyr3 and TyrCg (Table 5). These strains carried pSEVAb22-curs-dcs and pSEVAb83-comt-ccoaomttal-c3h plasmids.

**Table 5:** Production of curcumin from tyrosine (TYR). The mean and standard deviation of three replicates are shown. The yield is expressed as a percentage of the maximum theoretical yield. p22 and p83 refer to pSEVAb22 and pSEVAb83 plasmid backbones, respectively.

| Name | Plasmids | TYR (mM) | Titer (mM) | Yield (%) |
| --- | --- | --- | --- | --- |
| **Tyr1** | p22-*curs1-dcs* + p83-*comt-ccoaomt-tal-c3h* | 1 | 0.01 ± 0.00 | 1.30 ± 0.01 |
| | | 3 | 0.00 ± 0.00 | 0.31 ± 0.31 |
| **Tyr2** | p22-*curs2-dcs* + p83-*comt-ccoaomt-tal-c3h* | 1 | 0.01 ± 0.00 | 2.67 ± 0.21 |
| | | 3 | 0.01 ± 0.00 | 0.71 ± 0.21 |
| **Tyr3** | p22-*curs3-dcs* + p83-*comt-ccoaomt-tal-c3h* | 3 | 0.00 ± 0.00 | 0.00 ± 0.00 |
| **TyrCg** | p22-*cursCg-dcs* + p83-*comt-ccoaomt-tal-c3h* | 1 | 0.01 ± 0.00 | 1.11 ± 0.16 |
| | | 3 | 0.00 ± 0.00 | 0.10 ± 0.17 |

Considering the low tyrosine toxicity observed in the toxicity assay and the expected lower curcuminoid production from this substrate^9–11,28^, 3 mM tyrosine was used as initial substrate concentration. However, low curcuminoids concentrations were found for Tyr1, Tyr2 and TyrCg and no production was observed for Tyr3 (Table 5, Sup. Figure 5). Therefore, a new experiment with 1 mM of tyrosine was performed using the producing strains Tyr1, Tyr2 and TyrCg. Reducing the concentration of tyrosine reduced the lag phase of the cultures and improved curcuminoids production (Figure 1A, Table 5, Sup. Figure 5). The highest production was achieved by Tyr2 (0.013 ± 0.001 mM) followed by Tyr1 (0.010 ± 0.005) and TyrCg (0.005 ± 0.001). Notably, although production of all curcuminoids was possible, curcumin was the only curcuminoid measured. The highest curcumin yield achieved (2.7 ± 0.2% of the maximum theoretical yield) was higher than previously reported yields achieved with *E. coli*^9^. As observed with the other substrates, curcumin concentration decreased after 70 h indicating the degradation of this compound (Sup. Figure 5).

Additionally, Tyr1, Tyr2, Tyr3 and TyrCg were grown with glucose as only substrate to study curcuminoids production from endogenous tyrosine levels. Although no curcuminoids were detected in the Tyr3 strain, low curcumin concentrations were detected in cultures with Tyr1, Tyr2 and TyrCg (Sup. Figure 5). Even though the achieved concentrations were low, curcuminoids production from glucose by *P. putida* Δ*ech* expressing the complete curcuminoid pathway was demonstrated.

### Model-based optimization of curcumin production from tyrosine

Although production of curcuminoids from ferulic acid, p-coumaric acid, caffeic acid and tyrosine was achieved, maximum theoretical yields were not obtained (Figure 1), indicating the possibility of further pathway optimization. To facilitate this optimization, we developed kinetic models of the curcuminoid pathway, allowing to monitor the concentrations of biomass, substrate, products, and intermediates over time. These models enable the calculation of reaction fluxes and the impact of enzyme levels on production. Consequently, the insights gained through simulations were leveraged to steer the pathway optimization process. Considering that the lowest measured production was achieved with tyrosine as substrate and that curcumin was the main curcuminoid found, the optimization strategy focused on improving curcumin production from tyrosine.

#### Construction of kinetic models of the curcuminoids pathway

Kinetic models of the curcuminoid pathways expressed in FeCua2, Caa2 and Tyr2 strains were created. Considering that these models shared some reactions and their corresponding parameters (Table 2, Figure 2A-C), experimental data obtained with all these strains was simultaneously used for the parameterization of the models. Although this approach aimed to facilitate parameter estimation including experiments performed with different strains and substrates, it was not sufficient to obtain accurate parameter estimates. Instead, ensemble models formed by the 10 estimated parameter sets with the best agreement to experimental data (lowest least squares) were created and used for simulations (enFeCua2, enCaa2, enTyr2).

Figure 2D shows the fit of the ensemble models to the experiments used during parameter estimation. All models accurately described precursor profiles except enFeCua2 that incorrectly simulated ferulic acid accumulation with ferulic acid as substrate. All models accurately simulated curcumin and bisdemethoxycurcumin production with normalized mean square errors (nMSE) of 2.2% and 3.5%. Demethoxycurcumin could only be produced by the Tyr2 strain, used in one of the four experiments for model training, which resulted in a higher nMSE (23.2%) when simulating the production of this metabolite. Besides, model simulations pointed at accumulation of feruloyl-CoA, coumaroyl-CoA and, to a lesser extent, caffeoil-CoA and diketide forms as cause for curcuminoid yields below 100 % (Sup. Figure 6).

The ensemble modeling approach was validated simulating the performance of the FeCua2 strain simultaneously using ferulic and p-coumaric acids as substrates. Model predictions of curcuminoids showed good agreement with experimental data with a nMSE of 1.9%, 8.6% and 1.2% for CUR, BDCUR and DCUR, respectively (Figure 2E). Even though the ensemble failed to predict the complete depletion of ferulic acid, it predicted curcumin as main curcuminoid, capturing the reported preference of CURS2 towards feruloyl-CoA^37^.

#### Alleviation of enzymatic bottlenecks

After validating the qualitative performance of the ensemble models, the focus was set on optimizing curcumin production from tyrosine. Therefore, the enTyr2 ensemble model was used to calculate the maximum fluxes through each curcuminoid pathway reaction during production with tyrosine as substrate (Figure 3I). Fluxes through the last two steps of the pathway (DCS and CURS) and the FCS reaction consuming ferulic acid were lower than those from the upper reactions (TAL, C3H, COMT, CCOAOMT), which explained the accumulation of pathway intermediates (Sup. Figure 6). In the Tyr2 strain, *curs2* and *dcs* genes were expressed in a plasmid with a lower copy number compared to the other genes in the pathway (Figure 3A). The expected lower concentration of CURS2 and DCS enzymes was consistent with the lower fluxes calculated by the ensemble.

The flux imbalance among pathway reactions was addressed in the Tyr2 Opt1 and Tyr2 Opt2 strains by introducing *curs2* and *dcs* in higher copy number plasmids. While Tyr2 Opt1 carried *curs2* and *dcs* in the high-copy number backbone pSEVAb25 and *comt, ccoaomt, tal* and *c3h* in the medium-copy number backbone pSEVAb83; Tyr2 Opt2 carried these genes in the medium-copy number plasmid pSEVAb83 and the low-copy number plasmid pSEVAb22 (Table 6, Figure 3B, C). Production experiments were performed using 1 mM tyrosine as substrate and resulted in decreased production of curcumin by Tyr2 Opt1 and increased production of Tyr2 Opt2 (Figure 3J). While Tyr2 only produced curcumin at a final concentration of 0.013 ± 0.001 mM, Tyr2 Opt2 produced 0.021 ± 0.003mM curcumin and 0.031 mM of total curcuminoids (Figure 3J). This represented a 1.6-fold increase in curcumin production, 2.4-fold increase in curcuminoids production and a curcuminoids yield of 5.7 ± 0.3 % of the theoretical maximum. The effect of changing plasmid copy number was also assessed in strains expressing *curs1* codon-optimized for *C. glutamicum* instead of *curs2*, and similar results were found (Sup. Data 2).

**Table 6:** Production of curcumin from tyrosine (TYR) of optimized strains. The mean and standard deviation of three replicates are shown. The yield is expressed as a percentage of the maximum theoretical yield. p22, p83, and p25 refer to pSEVAb22, pSEVAb83, and pSEVAb25 plasmid backbones, respectively.

| Name | Plasmids | TYR (mM) | Titer (mM) | Yield (%) |
| --- | --- | --- | --- | --- |
| **Tyr2_Opt1** | p25-*curs2-dcs* + p83-*comt-ccoamt-tal-c3h* | 1 | 0.01 ± 0.00 | 1.67 ± 0.13 |
| **Tyr2_Opt2** | p83-*curs2-dcs* + p22-*comt-ccoamt-tal-c3h* | 1 | 0.02 ± 0.00 | 4.19 ± 0.58 |
|  |  | 2 | 0.06 ± 0.00 | 5.68 ± 0.33 |
| **Tyr2_Opt3** | p83-*curs2-dcs-c3h* + p22-*comt-ccoamt-tal* | 2 | 0.07 ± 0.01 | 6.97 ± 0.46 |
| **Tyr2_Opt4** | p83-*curs2-dcs* + p22-*ccoamt-tal-c3h* | 2 | 0.03 ± 0.01 | 3.13 ± 0.66 |
| **Tyr_Opt5** | p83-*curs2-dcs* + p22-*comt-ccoamt-tal-c3h-4cl4* | 2 | 0.08 ± 0.01 | 8.22 ± 0.57 |
| **Tyr2_Opt6** | p83-*curs2-dcs-4cl4* + p22-*comt-ccoamt-tal-c3h* | 2 | 0.03 ± 0.00 | 3.07 ± 0.30 |
| **Tyr2_Opt7** | p83-*curs2-dcs-c3h* + p22-*comt-ccoamt-tal-4cl4* | 2 | 0.11 ± 0.02 | 10.82 ± 1.81 |

Enhanced curcuminoid production was achieved by increasing the copy number of *dcs* and *curs2*. However, this improvement was only observed when coupled with a reduced expression of the other pathways genes, underscoring that overall higher gene expression does not guarantee a higher production. Rather, carefully selecting the genes to over-express was required. To facilitate this task, we employed the enTyr2 ensemble model to systematically evaluate the impact of varying enzyme concentrations on curcumin production from tyrosine. For each enzyme in the pathway, the effect of reducing and increasing its concentration was assessed simulating curcumin production with the enzyme concentration estimated for each model in the ensemble as reference (Sup. Figure 7).

Increasing the concentration of C3H and decreasing the concentration of COMT had the biggest positive impact on curcumin concentration in silico (Figure 3K). While increasing the concentration of C3H facilitates the consumption of p-coumaric acid; decreasing the concentration of COMT limits the production of ferulic acid which is predicted to accumulate due to the lower consumption flux by FCS (Figure 3I). In order to test model predictions, strains Tyr2 Opt3, with higher expression of C3H, and Tyr2 Opt4, not expressing COMT, were constructed and tested with 2 mM of tyrosine (Table 6, Figure 3D, E). Tyr2 Opt3 produced 0.07 ± 0.01 mM of curcumin, improving curcumin production 1.3-fold compared to Tyr2 Opt2. However Tyr2 Opt4 produced 24.4% less curcumin than Tyr2 Opt2 (Figure 3L). The decreased production observed in Tyr2 Opt4 was accompanied by the accumulation of caffeic and ferulic acids, which suggested the conversion of feruloyl-CoA to ferulic acid by FCS, a behavior only predicted by one of the ten models of the ensemble (Sup. Figure 8).

#### Expression of 4CL4 improves curcumin production

FCS was the only endogenous enzyme used in the curcuminoids pathway. Although this enzyme can convert p-coumaric, caffeic and ferulic acid into their CoA-forms, kinetic information regarding substrate preference was not available. Besides, endogenous regulation of *fcs* expression in response to metabolites in the curcuminoid pathway is unknown. Although FCS related parameters could not be estimated, model simulations and experimental data suggested a possible detoxifying role of FCS that could result in the conversion of caffeic acid into ferulic acid which negatively impacted curcuminoids production. To reduce the accumulation of ferulic acid, alternatives to *fcs* were sought. *C. longa* and *Arabidopsis thaliana* express 4CL enzymes that perform function analogous to FCS. Considering the kinetic parameters of these enzymes, expression of 4CL4 from *A. thaliana* was selected due to the 7.6-fold lower km of ferulic acid compared to 4CL1 from *C. longa*^*44*^.

The role of 4CL4 to complement FCS functions in vivo was tested in three strains: Tyr2 Opt5, Tyr2 Opt6 and Tyr2 Opt7 (Table 6, Figure 3F-H). Tyr2 Opt5 and Tyr2 Opt6 were based on Tyr2 Opt2 and were used to test the effect of expressing *4cl4* in pSEVAb22 and pSEVAb83 respectively. While expression of *4cl4* in the low-copy number plasmid improved curcumin production up to 0.08 ± 0.01 mM, expression in the medium-copy number plasmid reduced production to 0.03 ± 0.00 mM. When the expression of *4cl4* in pSEVAb22 was combined with the expression of *c3h* in pSEVAb83 in the Tyr2 Opt7 strain, 0.11 ± 0.02 mM of curcumin were achieved, a 1.6-fold increase compared to Tyr2 Opt3 and 4-fold increase compared to Tyr2 (Figure 3L).

## Discussion

In this study, we demonstrated the potential of using *P. putida* Δ*ech* as a platform for the bio-based production of curcuminoids, especially curcumin. We followed a systematic metabolic engineering framework, progressively expanding the substrate repertoire for curcuminoid production. Each step introduced new genes, increasing the pathway’s complexity. Additionally, we evaluated, for the first time, the impact of different *curs* isoenzymes on production, achieving the highest reported conversion of caffeic acid and tyrosine to curcumin when utilizing *cursCg* and *curs2*, respectively. The acquired data confirmed the importance of the pathway kinetics on production and served as the foundation for training kinetic models of the curcuminoid pathway. These models enhanced the interpretation of the conducted experiments, proposed the development of new strains, and guided the hypothesis generation process during pathway optimization. This methodology culminated in a 4-fold enhancement in tyrosine conversion to curcumin, increasing the yield from 2.7 ± 0.2 % to 10.8 ± 1.8 % of the theoretical maximum, a 8.5-fold improvement over previously reported conversions^9^.

Although CURS1, CURS2 and CURS3 have a high amino acid identity, their kinetic parameters are considerably different^37^. While CURS1 has the lowest affinity (KM) for ferulic and p-coumaric acid-derived molecules, it has the highest turnover rate (Kcat). In contrast, CURS3 shows the highest affinity towards these substrates but the lowest turnover rate. CURS2 takes a middle position, and possibly the combination of high affinity with twice the turnover rate of CURS3 is responsible for the better conversions shown in this study. Moreover, the preference of the FeCua2 strain for the ferulic acid conversion over p-coumaric acid conversion, is also explained by the 20-fold lower KM constant of this enzyme to feruloyl-CoA compared to coumaroyl-CoA^37^. Notably, despite the consistent accumulation of ferulic acid when caffeic acid is used as a substrate, strains expressing curs1 codon-optimized for *P. putida* or *C. glutamicum* achieved higher conversion rates compared to *strains* expressing *curs2*. A higher toxicity of caffeic acid on the Caa2 strain was discarded as the cause of the lower conversion since the growth of this strain was comparable to the Caa1 and CaaCg strains.

The significance of the kinetic parameters of CURS enzymes in production became apparent, emphasizing the critical role of kinetics in optimizing pathway performance. This understanding was crucial given the promiscuity not only of CURS but also of FCS and DCS enzymes. Consequently, we constructed kinetic models of the pathway. Although accurate parameter estimation was not achieved, we developed ensemble models for the FeCua2, Caa2 and Tyr2 strains. Flux analyses using the enTyr2 model, combined with simulations assessing the influence of enzyme concentration on production, guided the development of the Tyr2 Opt3 strain. Within this strain, the genes *dcs, curs2* and *c3h* are expressed in the medium-copy number plasmid (pSEVAb83) while *comt, ccoaomt* and *tal* are expressed in a low-copy number plasmid (pSEVAb22). This strain achieved a 7.0 ± 0.5 % conversion of tyrosine to curcumin, a 2.6-fold improvement compared to Tyr2.

Despite the consistent accuracy of ensemble models in simulating curcuminoid production, they were unable to replicate the complete depletion of ferulic acid in the absence of caffeic acid or tyrosine. This disagreement between experimental data and simulations was key to identifying the need to complement the endogenous expression of *fcs* with the expression of *A. thaliana 4cl4*. Maximum reaction fluxes calculated by enTyr2 revealed a low flux of the FCS reaction consuming ferulic acid compared to the FCS reactions consuming p-coumaric and caffeic acids (Figure 3I). Besides, model simulations suggested omitting the expression of *comt* to improve curcumin production (Figure 3K). These observations led to the hypothesis that, in the absence of caffeic acid, FCS is able to convert ferulic acid into its CoA form but, when caffeic acid is present, FCS saturates with p-coumaric and caffeic acids resulting in the accumulation of ferulic acid. This accumulation is accentuated by the direct conversion of caffeic acid into coumaric acid by COMT. Although model simulations suggested reducing *comt* expression to improve curcumin production, excluding this gene in the Tyr2 Opt4 strain decreased curcumin production and led to ferulic acid accumulation (Figure 3L). In this strain, FCS catalyzes the reverse conversion from feruloyl-CoA to ferulic acid, a mechanism only captured by one of the models in the ensemble. Therefore, we addressed the possible limitation caused by FCS by expressing *4cl4* in the Tyr2 Opt7 strain. As a result, we achieved a conversion efficiency of 10.8 ± 1.8 % from tyrosine to curcumin, 1.6-fold and 4-fold improvement compared to Tyr2 Opt3 and Tyr2, respectively. In addition, we showed the ability of the ensemble modeling approach to study the curcuminoid pathway even when accurate parameter values cannot be estimated.

Beyond gene expression, the observed degradation of curcumin, the substrate concentration, and the initial OD600nm of the culture influenced the curcumin yields. Photodegradation of curcuminoids is well-documented^1^ and can be prevented by shielding the cultivation medium from light exposure. Next to the (photo)chemical instability, the NADPH-dependent curcumin/dihydrocurcumin reductase (*curA* gene) can catalyze the breakdown of curcumin. The deletion of this gene improved curcumin yield in *E. coli*^*23*^ and could be implemented in *P. putida* Δ*ech*. Furthermore, the initial substrate concentration was found to be a major determinant in curcuminoid production outcomes. The optimum substrate concentration varied depending on the strain, with strains exhibiting superior conversion capabilities tolerating higher substrate concentrations. Including the effect of substrate and intermediate concentrations on growth in the ensemble models could therefore be used to estimate optimum initial concentrations and further enhance production. Alongside substrate concentration, other processrelated parameters can be tuned to improve production^20^. For example, we showed how increasing the initial OD600nm of the cultures resulted in a 0.7-fold increase in curcumin production from ferulic acid (Table 3, Sup. Figure 2).

Remarkably, *P. putida* Δ*ech* showed lower conversion yields when using ferulic and p-coumaric acids as substrates compared to *E. coli*^*9,11*^. However, as the complexity of the pathway increased, and caffeic acid and tyrosine were used as substrates, *P. putida* Δ*ech* strains out-competed previously reported *E. coli* strains^9,11^. The 10-fold increase in caffeic acid conversion by CaaCg, Caa1 and Caa2 compared to *E. coli* could be attributed to the expression of *comt* or the higher tolerance of *P. putida* to this substrate. Although conversions up to 11.5 % have been reported with tyrosine as substrate, they resulted in the production of bisdemethoxycurcumin^10,37^. These strains were unable to produce caffeic and ferulic acids (Table 1) which reduced the number of possible toxic pathway metabolites and likely favored production. When *c3h* and *comt* were expressed in one of these strains, the production of curcumin was reduced to 0.1 % of the maximum theoretical yield^28^. This yield was improved by Rodrigues et al. to 1.3 % expressing *dcs* and *curs1* instead of *cus*^*9*^, probably due to the higher affinity of *curs1* to ferulic acid-derived molecules compared to *cus*^*12*^. Higher yields of curcumin from tyrosine (2.9% of the theoretical maximum) have only been achieved by the use of a co-culture with two *E. coli* strains, each expressing a part of the pathway^9^. Notably, the Tyr2 strain alone matched the yield achieved with the co-culture, and Tyr2 Opt7 showed an 8.5-fold improvement, which highlights the potential of *P. putida* for curcumin production.

In summary, we establish the production of curcuminoids in *P. putida*, optimize production yields from ferulic acid, p-coumaric acid, caffeic acid, and tyrosine, and provide the basis to produce curcuminoids from glucose. The study’s strength lies in its systematic approach: utilizing ensemble dynamic models to understand pathway kinetics and identify bottlenecks, testing various isoenzymes for their efficiency, and fine-tuning gene expression levels and substrate concentrations for optimal yields. This comprehensive approach led to significant improvements in curcuminoid production, culminating in the highest yield (10.8 ± 1.8 % of the theoretical maximum) of curcumin production from tyrosine reported to date.

## References

1. Nelson, K. M. et al. The Essential Medicinal Chemistry of Curcumin. J. Med. Chem. 60, 1620–1637 (2017).

2. Regulation (EC) No 1333/2008 of the European Parliament and of the Council of 16 December 2008 on food additives. 16–33 (2008).

3. Listing of color additives exempt from certification. (Food and Drug Administration, 2001).

4. Research, G. V. Curcumin Market Size, Share & Trends Analysis Report By Application (Pharmaceutical, Food, Cosmetics), By Region (North America, Europe, Asia Pacific, CSA, MEA), And Segment Forecasts, 2020–2028. (2018).

5. Heger, M. Don’t discount all curcumin trial data. Nat. 2017 5437643 543, 40–40 (2017).

6. Iturbide, M. et al. An update of IPCC climate reference regions for subcontinental analysis of climate model data: definition and aggregated datasets. Earth Syst. Sci. Data 12, 2959–2970 (2020).

7. Jiang, T., Ghosh, R. & Charcosset, C. Extraction, purification and applications of curcumin from plant materials-A comprehensive review. Trends Food Sci. Technol. 112, 419–430 (2021).

8. Indira Priyadarsini, K. molecules The Chemistry of Curcumin: From Extraction to Therapeutic Agent. Molecules 19, (2014).

9. Rodrigues, J. L., Gomes, D. & Rodrigues, L. R. A Combinatorial Approach to Optimize the Production of Curcuminoids From Tyrosine in Escherichia coli. Front. Bioeng. Biotechnol. 8, 509978 (2020).

10. Katsuyama, Y., Matsuzawa, M., Funa, N. & Horinouchi, S. Production of curcuminoids by Escherichia coli carrying an artificial biosynthesis pathway. Microbiology 154, 2620–2628 (2008).

11. Rodrigues, J. L., Araújo, R. G., Prather, K. L. J., Kluskens, L. D. & Rodrigues, L. R. Production of curcuminoids from tyrosine by a metabolically engineered Escherichia coli using caffeic acid as an intermediate. Biotechnol. J. 10, 599–609 (2015).

12. Katsuyama, Y., Matsuzawa, M., Funa, N. & Horinouchi, S. In vitro synthesis of curcuminoids by type III polyketide synthase from Oryza sativa. J. Biol. Chem. 282, 37702–37709 (2007).

13. Mishra, S., Wang, Z., Volk, M. J. & Zhao, H. Design and application of a kinetic model of lipid metabolism in Saccharomyces cerevisiae. Metab. Eng. 75, 12–18 (2023).

14. Martin, J. et al. A dynamic kinetic model captures cell-free metabolism for improved butanol production. Metab. Eng. (2023) doi:10.1016/J.YMBEN.2023.01.009.

15. Nikel, P. I., Chavarría, M., Chavarría, M. & de Lorenzo, V. From dirt to industrial applications: Pseudomonas putida as a Synthetic Biology chassis for hosting harsh biochemical reactions. Curr. Opin. Chem. Biol. 34, 20–29 (2016).

16. Martin-Pascual, M. et al. A navigation guide of synthetic biology tools for Pseudomonas putida. Biotechnol. Adv. 49, 107732 (2021).

17. Poblete-Castro, I. et al. The metabolic response of P. putida KT2442 producing high levels of polyhydroxyalkanoate under single-and multiple-nutrient-limited growth: Highlights from a multi-level omics approach. (2012).

18. Loeschcke, A. & Thies, S. Pseudomonas putida—a versatile host for the production of natural products. Appl. Microbiol. Biotechnol. 99, 6197–6214 (2015).

19. Senior, A. W. et al. Improved protein structure prediction using potentials from deep learning. Nat. 2020 5777792 577, 706–710 (2020).

20. Couto, M. R., Rodrigues, J. L. & Rodrigues, L. R. Optimization of fermentation conditions for the production of curcumin by engineered Escherichia coli. J. R. Soc. Interface 14, (2017).

21. Rodrigues, J. L. et al. Hydroxycinnamic acids and curcumin production in engineered Escherichia coli using heat shock promoters. Biochem. Eng. J. 125, 41–49 (2017).

22. Ciurkot, K., Gorochowski, T. E., Roubos, J. A. & Verwaal, R. Efficient multiplexed gene regulation in Saccharomyces cerevisiae using dCas12a. Nucleic Acids Res. 49, 7775–7790 (2021).

23. Wu, Y. et al. Design of a programmable biosensor-CRISPRi genetic circuits for dynamic and autonomous dual-control of metabolic flux in Bacillus subtilis. Nucleic Acids Res. 48, 996–1009 (2020).

24. Rainha, J., Rodrigues, J. L., Faria, C. & Rodrigues, L. R. Curcumin biosynthesis from ferulic acid by engineered Saccharomyces cerevisiae. Biotechnol. J. 17, 2100400 (2022).

25. Fang, Z., Jones, J. A., Zhou, J. & Koffas, M. A. G. Engineering Escherichia coli Co-Cultures for Production of Curcuminoids From Glucose. Biotechnol. J. 13, 1700576 (2018).

26. Incha, M. R. et al. Leveraging host metabolism for bisdemethoxycurcumin production in Pseudomonas putida. Metab. Eng. Commun. 10, e00119 (2020).

27. Palmer, C. M., Miller, K. K., Nguyen, A. & Alper, H. S. Engineering 4-coumaroyl-CoA derived polyketide production in Yarrowia lipolytica through a β-oxidation mediated strategy. Metab. Eng. 57, 174–181 (2020).

28. Wang, S. et al. Metabolic engineering of Escherichia coli for the biosynthesis of various phenylpropanoid derivatives. Metab. Eng. 29, 153–159 (2015).

29. Green, R. & Rogers, E. J. Transformation of Chemically Competent E. coli. Methods Enzymol. 529, 329–336 (2013).

30. Volke, D. C., Friis, L., Wirth, N. T., Turlin, J. & Nikel, P. I. Synthetic control of plasmid replication enables target- and self-curing of vectors and expedites genome engineering of Pseudomonas putida. Metab. Eng. Commun. 10, e00126 (2020).

31. Damalas, S. G., Batianis, C., Martin-Pascual, M., de Lorenzo, V. & Martins dos Santos, V. A. P. SEVA : enabling interoperability of DNA assembly among the SEVA, BioBricks and Type IIS restriction enzyme standards. Microb. Biotechnol. 13, 1793–1806 (2020).

32. Liebermeister, W., Uhlendorf, J. & Klipp, E. Modular rate laws for enzymatic reactions: thermodynamics, elasticities and implementation. Bioinformatics 26, 1528–1534 (2010).

33. Gläser, L. et al. A common approach for absolute quantification of short chain CoA thioesters in prokaryotic and eukaryotic microbes. Microb. Cell Fact. 19, 1–13 (2020).

34. Wordofa, G. G. et al. Quantifying the Metabolome of Pseudomonas taiwanensis VLB120: Evaluation of Hot and Cold Combined Quenching/Extraction Approaches. Anal. Chem. 89, 8738–8747 (2017).

35. Bennett, B. D. et al. Absolute metabolite concentrations and implied enzyme active site occupancy in Escherichia coli. Nat. Chem. Biol. 5, 593–9 (2009).

36. Noor, E., Haraldsdóttir, H. S., Milo, R. & Fleming, R. M. T. Consistent Estimation of Gibbs Energy Using Component Contributions. PLOS Comput. Biol. 9, e1003098 (2013).

37. Katsuyama, Y., Kita, T. & Horinouchi, S. Identification and characterization of multiple curcumin synthases from the herb Curcuma longa. (2009) doi:10.1016/j.febslet.2009.07.029.

38. Katsuyama, Y., Miyazono, K. I., Tanokura, M., Ohnishi, Y. & Horinouchi, S. Structural and biochemical elucidation of mechanism for decarboxylative condensation of β-keto acid by curcumin synthase. J. Biol. Chem. 286, 6659–6668 (2011).

39. Lubitz, T., Schulz, M., Klipp, E. & Liebermeister, W. Parameter balancing in kinetic models of cell metabolism. J. Phys. Chem. B 114, 16298–16303 (2010).

40. Beber, M. E. et al. eQuilibrator 3.0: a database solution for thermodynamic constant estimation. Nucleic Acids Res. 50, D603–D609 (2022).

41. Stapor, P., Fröhlich, F. & Hasenauer, J. Optimization and profile calculation of ODE models using second order adjoint sensitivity analysis. Bioinformatics 34, i151–i159 (2018).

42. Fröhlich, F., Kaltenbacher, B., Theis, F. J. & Hasenauer, J. Scalable Parameter Estimation for Genome-Scale Biochemical Reaction Networks. PLOS Comput. Biol. 13, e1005331 (2017).

43. von Kamp, A. & Klamt, S. Enumeration of Smallest Intervention Strategies in Genome-Scale Metabolic Networks. PLoS Comput. Biol. 10, e1003378 (2014).

44. Hamberger, B. & Hahlbrock, K. The 4-coumarate:CoA ligase gene family in Arabidopsis thaliana comprises one rare, sinapate-activating and three commonly occurring isoenzymes. Proc. Natl. Acad. Sci. 101, 2209–2214 (2004).

